# New de novo assembly of the Atlantic bottlenose dolphin (*Tursiops truncatus*) improves genome completeness and provides haplotype phasing

**DOI:** 10.1101/376301

**Authors:** Karine A. Martinez-Viaud, Cindy Taylor Lawley, Milmer Martinez Vergara, Gil Ben-Zvi, Tammy Biniashvili, Kobi Baruch, Judy St. Leger, Jennie Le, Aparna Natarajan, Marlem Rivera, Marbie Guillergan, Erich Jaeger, Brian Steffy, Aleksey Zimin

## Abstract

High quality genomes are essential to resolve challenges in breeding, comparative biology, medicine and conservation planning. New library preparation techniques along with better assembly algorithms result in continued improvements in assemblies for non-model organisms, moving them toward reference quality genomes. We report on the latest genome assembly of the Atlantic bottlenose dolphin leveraging Illumina sequencing data coupled with a combination of several library preparation techniques. These include Linked-Reads (Chromium, 10x Genomics), mate pairs, long insert paired ends and standard paired ends. Data were assembled with the commercial DeNovoMAGIC^TM^ assembly software resulting in two assemblies, a traditional “haploid” assembly (Tur_tru_Illumina_hap_v1) that is a mosaic of the two parental haplotypes and a phased assembly (Tur_tru_Illumina_phased_v1) where each scaffold has sequence from a single homologous chromosome. We show that Tur_tru_Illumina_hap_v1 is more complete and accurate compared to the current best reference based on the amount and composition of sequence, the consistency of the mate pair alignments to the assembled scaffolds, and on the analysis of conserved single-copy mammalian orthologs. The phased de novo assembly Tur_tru_Illumina_phased_v1 is the first publicly available for this species and provides the community with novel and accurate ways to explore the heterozygous nature of the dolphin genome.

## Introduction

Technical advances in the past decade have reduced sequencing costs and improved access to sequencing data. Subsequent improvements in DNA extraction, preparation, and assembly algorithms facilitate low cost accurate de novo genome assemblies. Such assemblies are essential for constructing haplotype diversity databases for breeding, comparative biology, medicine and conservation planning. Even highly complex genomes now benefit from higher contiguity and improved protein coding coverage [1-4]. Consortium efforts to catalogue biodiversity of pivotal species of comparative evolutionary significance will continue to drive novel low-cost approaches toward reference quality assemblies with chromosome level resolution [5-7]. Here we use a combination of methods to drive improvements in assembly structure for the Atlantic bottlenose dolphin (*Tursiops truncatus*). This genome assembly, like that of the Hawaiian Monk seal and African wild dog, is being published with the goal to facilitate research on comparative genomics, provide structure for cataloging biodiversity and ultimately support decisions around species conservation and management [8, 9].

The bottlenose dolphin is one of the most widely studied marine mammals, however the taxonomy of the *Tursiops* genus remains unresolved. Numerous species designations have been suggested but not adopted due to a lack of resolution afforded by available data [10]. Even with new molecular genetic markers, we have reached a limitation on resolution from genetic data available to delineate species, subspecies and populations [11]. To usher this species into the era of genomics, a high-quality reference genome is essential. It provides structure to catalogue diversity within and between species at the whole genome level. In addition, the parallel molecular trajectory between dolphin and other mammalian species [12] makes the bottlenose dolphin a useful model to understand aspects of human health such as metabolic processes/diabetes [13-15], proteomics [16, 17] and aging [18].

A preliminary dolphin genome was first submitted to NCBI (TurTru1.0; GCA_000151865.1) using low coverage (2.82X) Sanger sequencing for the purpose of cross-species comparison [12, 19, 20]. Subsequent improvements were achieved through the addition of 30X Illumina short read data and 3.5X 454 data (Ttru_1.4; GCA_000151865.3). A much more complete genome was submitted in 2016 leveraging improvements in library preparation and assembly methods (Meraculous v. 2.2.2.5 and HiRise v. 1.3.0-116-gf50c3ce; Dovetail, Inc) with 114X coverage of Illumina HiSeq data prepared with proximity ligation Hi-C protocol (Tur_tru v1; GCA_001922835.1; [16]).

With the collection of data from multiple sources including Linked-Reads (Chromium, 10x Genomics; [21]), mate pairs (MP), long insert paired ends and standard paired ends, and the DeNovoMAGIC assembly tool (NRGene, Ness-Ziona, Israel), we provide an improved haploid reference quality dolphin genome assembly as well as the first haplotype phased diploid assembly. We refer to our unphased assembly as Tur_tru_Illumina_hap_v1 and to the phased assembly as Tur_tru_Illumina_phased_v1. Using Tur_tru v1 for comparison, our assembly shows increased contiguity and completeness with high consistency to the MP data and orthologous mammalian protein alignments. Additionally, by aligning Tur_tru_Illumina_hap_v1 to the Human reference genome, we illustrate the synteny of the dolphin scaffolds to human chromosome 1 [22, 23].

## Results

### Coverage

We generated sequence data for a total coverage of approximately 450X, the majority from PCR Free and Chromium 10X Genomics Linked-Read libraries (Table 1). Coverage was computed using 2.4Gbp estimated genome size. Genome assembly was conducted using DeNovoMAGIC^TM^ software (NRGene, Ness-Ziona, Israel). More detail about the library preparation and the assembly process are found in the Methods section.

**Table 1.** Summary of the sequencing data collected to create Tur_tru_Illumina_hap_v1 and Tur_tru_Illumina_phased_v1.

|  |  |  |  |
| --- | --- | --- | --- |
| Library type | Read Length | Insert Size | Genomic Coverage |
| PCR-free | 2x250bp | 450bp | 101x |
| PCR-free | 2x160bp | 800bp | 123x |
| Mate-Pair | 2x150bp | 2-4Kbp (peak 4.2Kbp) | 37x |
| Mate-Pair | 2x150bp | 5-7Kbp (peak 6.0Kbp) | 61x |
| Mate-Pair | 2x150bp | 8-10Kbp (peak 9.9Kbp) | 58x |
| 10X Chromium | 2x150bp | - | 70x |

### Haploid and diploid assemblies

We report on two assemblies in this manuscript, one traditional haploid consensus assembly Tur_tru_Illumina_hap_v1 that represents a mosaic of the maternal and paternal haplotypes, and the other haplotype-phased (i.e., diploid) assembly where each scaffold represents sequence corresponding to a single haplotype, Tur_tru_Illumina_phased_v1. The quantitative statistics for both assemblies are listed in Table 2. The phased or diploid genome assembly was made possible using Illumina sequencing data by leveraging the combination of library prep methods including Linked-Reads, is a significant advance and will provide the community with a powerful genomic tool for the downstream analysis in the context of the true heterozygous dolphin genome.

**Table 2.** Comparison of quantitative statistics for different assemblies of the bottlenose dolphin. The total sequence listed excludes Ns (ambiguous nucleotides). Ns were also squeezed out from the scaffolds for N50 computations. We used genome size of 2,383,130,043bp equal to the total amount of sequence in the scaffolds of the bigger haploid assembly, for comparison of the N50 contig and scaffold sizes between the two assemblies. The Tur_tru_Illumina_hap_v1 and Tur_tru v1 assemblies have comparable scaffold N50 sizes, and Tur_tru v1 has bigger contigs. The Tur_tru_Illumina_hap_v1 assembly has more sequence and our BUSCO analysis (Table 3) shows that it is likely more complete. The N50 comparisons to the haplotype-resolved Tur_tru_Illumina_phased_v1 assembly are shown for completeness, computed with 2x genome size (2*2,383,130,043 = 4,766,260,086bp).

|  |  |  |  |
| --- | --- | --- | --- |
|  | Tur_tru v1 | Tur_tru_Illumina_hap_v1 | Tur_tru_Illumina_phased_v1 |
| Total sequence | 2,120,283,832 | 2,383,130,043 | 4,678,362,582 |
| # of scaffolds | 2647 | 481 | 98,209 |
| Longest scaffold | 96,299,184 | 83,924,496 | 10,429,594 |
| Scaffold N50 | 23,564,561 | 26,997,441 | 777,432 |
| Scaffold L50 | 26 | 30 | 1,509 |
| # of contigs | 116,650 | 139,544 | 355,974 |
| Longest contig | 403,070 | 320,783 | 298,006 |
| Contig N50 | 37,749 | 30,985 | 25,997 |
| Contig L50 | 17,321 | 23,199 | 53,738 |
| GC content | 40.85 | 41.25 | 41.95 |

### Genome assembly comparison

Both assemblies were compared to the best available assembly Tur_tru v1 (NCBI accession GCA_001922835.1; [16]). We did not use the Ttru_1.4 assembly (NCBI accession GCA_000151865.3) because the contiguity statistics of the Ttru_1.4 are vastly inferior to the Tur_tru v1 with a contig N50 3 times smaller than Tur_tru v1 and scaffold N50 over 200 times smaller.

The statistics for the Tur_tru_Illumina_hap_v1 assembly show bigger scaffolds but slightly smaller contigs with about 13% more sequence in the scaffolds compared to Tur_tru v1 (Table 2). More sequence does not necessarily make for a better assembly considering that the extra sequence may be duplicated haplotypes or contaminants that do not belong to the original organism. To characterize the extra sequence, we first aligned the Tur_tru_Illumina_hap_v1 to the Tur_tru v1 assembly using the Nucmer aligner which is part of MUMmer4 package [24]. We used default settings for generating the alignments. We then analyzed the alignments using the dnadiff package included with MUMmer4. 87.5% of Tur_tru_Illumina_hap_v1 sequence aligned to 97.9% of Tur_tru v1. This shows that 12.5% of Tur_tru_Illumina_hap_v1 had no alignments to Tur_tru v1, while only 2.1% of Tur_tru v1 had no alignments to Tur_tru_Illumina_hap_v1. Therefore, there are 301Mbp of extra novel sequence in our new assembly Tur_tru_Illumina_hap_v1. We then used the BUSCO tool to show that the extra sequence is meaningful (Table 3). Tur_tru_Illumina_hap_v1 had 160 missing BUSCOs, compared to 270 missing in Tur_tru v1. The number of duplicated BUSCOs was higher by only 34 in our assembly compared to Tur_tru v1. This suggests that most of the extra sequence in Tur_tru_Illumina_hap_v1 is not contamination or redundant sequence, and likely contains useful coding information. There were 105 BUSCOS missing from both assemblies. We examined the locations of the 165 BUSCOs that are only found in the Tur_tru_Illumina_hap_v1 and all of them fully or partially aligned to locations in the sequences that were missing in Tur_tru v1 assembly. Figure 1 shows the Venn diagram of BUSCOs aligned to both assemblies, showing that there are 165 BUSCOS that are only present in Tur_tru_Illumina_hap_v1 and 55 that are only present in Tur_tru v1, with 3779 present in both assemblies. The haplotype-resolved assembly is more fragmented and it is missing 266 BUSCOs. As expected most of the complete BUSCOs that were found (3537) are duplicated 2227), since they are found in different haplotypes.

**Table 3.**
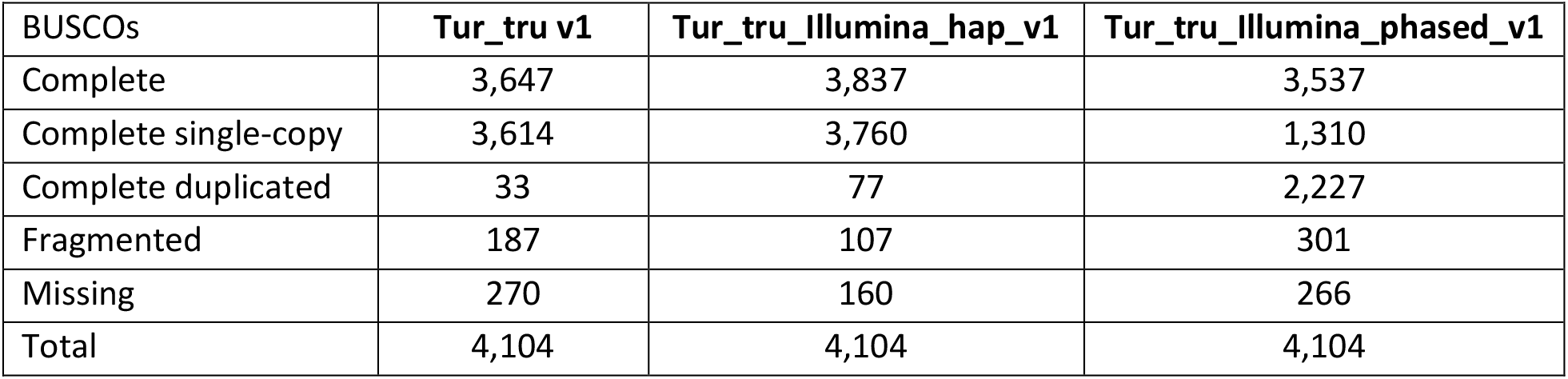
Comparison of BUSCO 3.0.2 Mammalia single copy ortholologs among the three Dolphin assemblies. The table shows that the Tur_tru_Illumina_hap_v1 assembly is more complete, with 110 fewer missing single-copy orthologs compared to the Tru_tru v1 assembly. The Tur_tru_Illumina_hap_v1 assembly has 43 extra duplicated orthologs, which possibly points to incomplete filtering of redundant haplotypes. While Tur_tru v1 assembly has bigger contigs, the Tur_tru_Illumina_hap_v1 assembly has many fewer fragmented BUSCOs. The haplotype-resolved Tur_tru_Illumina_phased_v1 assembly is less contiguous and less complete. As expected, more than half of the complete BUSCOs are duplicated, corresponding to the two resolved haplotypes.

**Figure 1.**
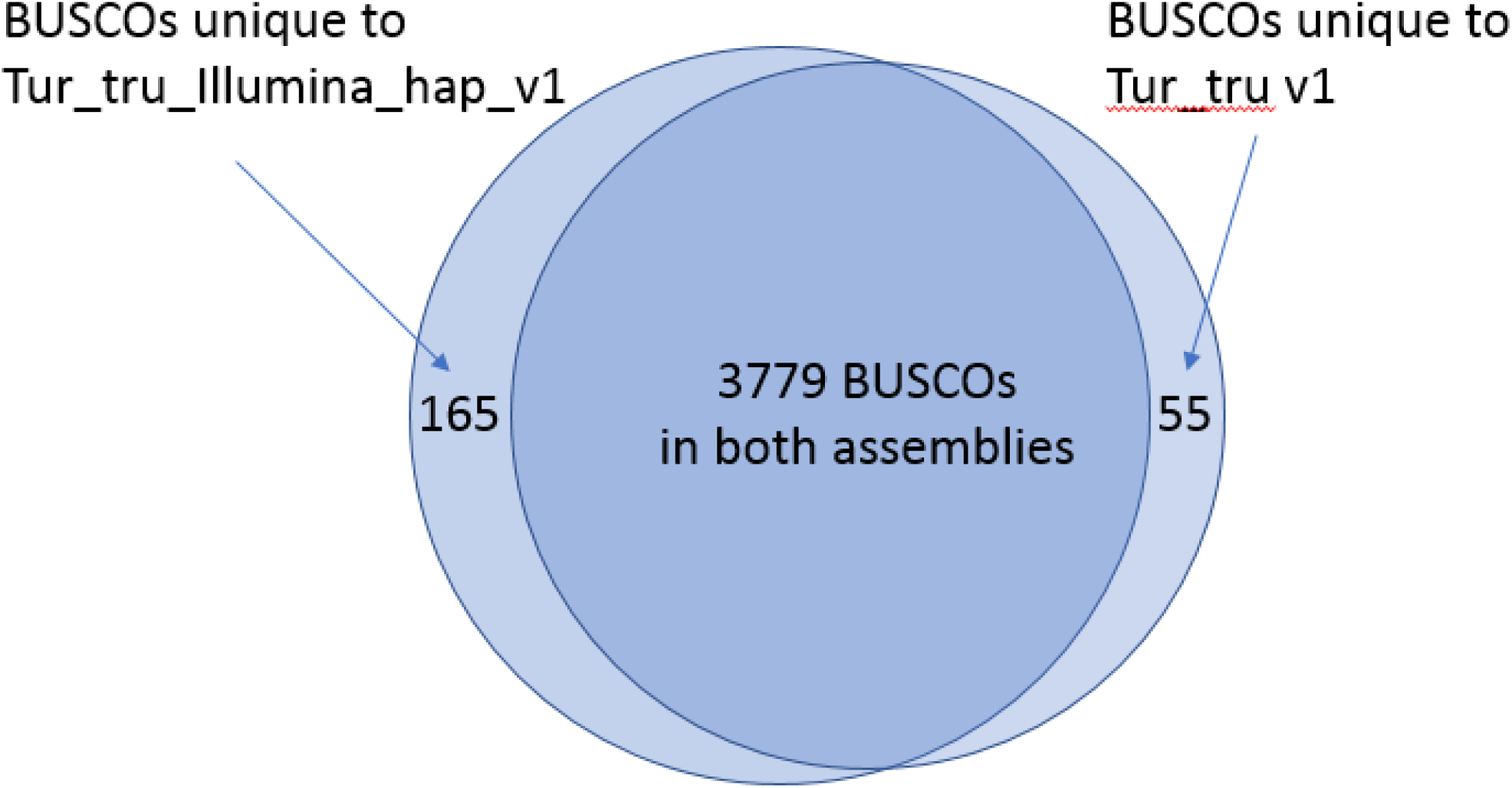
Venn diagram of BUSCOs present in two dolphin assemblies. Out of 4,104 BUSCOs in mammalia set, 105 are missing from both assemblies. Our assembly has 165 BUSCOs not present in Tur_tru v1 and Tur_tru v1 has 55 BUSCOs that are not present in our assembly.

### Assembly validation through MP consistency

Since both Tur_tru v1 and Tur_tru_Illumina_hap_v1 reference the same species, we expect few rearrangements between the assemblies. To examine this, we compared the absolute and relative correctness of the scaffolds of Tur_tru_Illumina_hap_v1 assembly by aligning the Illumina data from the 5-7Kbp MP library to the scaffolds of Tur_tru_Illumina_hap_v1, Tur_tru_Illumina_phased_v1,and Tur_tru v1 assemblies using the Bowtie2 tool [28]. We chose this library because it contained the largest number of valid 5-7Kbp mate pairs. We then used only high quality uniquely aligning mated reads (both mates had to align uniquely with quality score 42 in the SAM file) and classified the alignments of the MPs into the following categories (Table 4):

1. **Same scaffold happy** – number of MPs where both mates aligned to the same scaffold in the correct orientation with mate separation within 3 standard deviations of the library mean
2. **Same scaffold short** -- number of MPs where both mates aligned to the same scaffold in the opposite orientation with mate separation of less than 1000bp; these MPs are not indicative of scaffolding misassemblies, they are simply a byproduct of the mate pair library preparation process as they are MPs that are missing the circularization junction site between the mates
3. **Same scaffold long** – number of MPs where both mates aligned to the same scaffold in the correct orientation, but the mate separation exceeded three standard deviations of the library mean
4. **Same scaffold misoriented** -- number of MPs where both mates aligned to the same scaffold in the opposite orientation with mate separation of more than 1000bp
5. **Mates aligned to different scaffolds** – number of MPs where the two mates aligned to different scaffolds
6. **Only one mate in the pair aligned** – number of MPs where only one read aligned to the assembly.

**Table 4.** Comparison of the number of mate pairs (MPs) from 5-7Kbp library uniquely aligned to Tur_tru_Illumina_hap_v1, Tur_tru_Illumina_phased_v1 and Tur_tru v1 assemblies. The alignments were done with Bowtie2. Only the reads that mapped uniquely were used for this computation, thus the number of MPs uniquely mapping to haplotype resolved assembly is much smaller. Same scaffold means that both mates mapped to the same scaffold; happy mates aligned in the correct orientation with mate distance within 5 standard deviations from the mean; misoriented mates aligned in wrong orientation; long mates aligned with the distance between the mates exceeding 5 standard deviations; short mates aligned with the distance of less than 1000bp. Same scaffold mate ALL is the total number of all mate pairs where both mates aligned to the same scaffold.

|  | Tur_tru_Illumina_hap_v1 | Tur_tru_Illumina_phased_v1 | Tur_tru v1 |
| --- | --- | --- | --- |
| **Same scaffold mate ALL** | 228,285,253 | 73,514,546 | 219,277,963 |
| **Same scaffold happy** | 125,224,663 | 37,248,420 | 117,942,390 |
| **Same scaffold misoriented** | 158,547 | 51,533 | 1,164,118 |
| **Same scaffold long** | 82,948 | 9,183 | 274,458 |
| **Same scaffold short** | 102,819,095 | 36,205,410 | 99,896,997 |
| **Mates aligned to different scaffolds** | 5,629,393 | 7,438,471 | 5,576,802 |
| **One mate in the pair aligned** | 61,284,891 | 26,228,913 | 68,707,025 |

“Same scaffold ALL” category in table 4 is the sum of all mates in categories 1 to 4, it is listed for completeness.

Comparing Tur_tru_Illumina_hap_v1 with Tur_tru v1, the total number of reads uniquely aligning to both “haploid” assemblies is very similar: about 295.2M reads aligned to Tur_tru_Illumina_hap_v1 vs. about 293.6M reads aligned to Tur_tru v1. The total number of mate pairs aligning to the same scaffold is larger for Tur_tru_Illumina_hap_v1. Of the mate pairs aligning to the same scaffold, the number of mate pairs in the “Same scaffold happy” category is very similar between the two assemblies. The differences that stand out are the much larger (7.3 times more) number of mates that aligned to the same scaffold in the wrong orientation and the much larger (3.3 times more) number of the same scaffold long pairs in Tur_tru v1 compared to Tur_tru_Illumina_hap_v1 (Table 4). Of course, some level of discrepancy is expected, because the two assemblies represent two different individuals with unknown level of structural variation between them. However, in concert, the two different categories may also suggest a possibility of a relatively higher number of locally mis-ordered or misoriented contigs in the scaffolds of Tur_tru v1 assembly. This may be due to the scaffolding process used to create Tur_tru v1 assembly. The assembly was created with the HiRise assembler [25] using proximity ligation Hi-C data for scaffolding. The proximity ligation data provide MPs of all possible sizes, however, the MP distances and mate orientations are unknown. Since there are more shorter pairs than longer pairs due to the 3D structure of the DNA – it is much more likely to ligate parts of DNA that are closer to each other than the ones that are far apart. This property enables one to use these data for scaffolding. By mapping the pairs to the assembled scaffolds, one can measure how the distance between the mates in a pair varies with the number of pairs whose ends map to the same location in the assembly. However, the dependence is weak on the short end, meaning that the number of pairs of about 10Kbp in length is not much different from the number of links of 12-13Kbp in length. This frequently results in mis-orientations and shuffling of scaffold positions for contigs or scaffolds that are smaller than 10-20Kbp in the scaffolding process.

The haplotype phased assembly is much more fragmented, compared to both haploid assemblies, resulting in higher relative number of mate pairs mapping to different scaffolds. However, when looking at the “internal” mate pairs, i.e. where both mates map at least 10Kb away from the scaffold ends, we see remarkable consistency with less than 0.5% of the mates mapped to the wrong scaffold (see next section). Since for this analysis we only used mates mapping uniquely to the assembly, and there are two copies of the genome in the assembly, the total number of mapped mates is much lower.

### Haplotype resolution

To Illustrate the resolution of the haplotypes in Tur_tru_Illumina_phased_v1, we aligned it to Tur_tru_Illumina_hap_v1 using Nucmer tool. In Figure 2 we show the mummerplot of alignments of the phased assembly to the haploid one. The circles represent contig ends with lines joining them representing aligned sequence. The color indicates direction of the alignments. We display the alignments to an arbitrarily chosen scaffold314 of the haploid assembly. Only alignments longer than 5Kb are shown. Figure 2 shows that most of the “haploid” assembly aligns to two phased scaffolds, that is for each location on the x axis there are two corresponding alignments on the y axis. The regions that are covered by a single haplotype (rather than 2) are most probably homozygous regions of this genome. In cases where the homozygous region is long, it is more difficult to phase its heterozygous ends. Thus, in some cases, the homozygous regions are represented only once in the phased assembly (instead of twice). This is the cause for some of the single copy BUSCOs in the phased assembly. In our experience, this issue is more pronounced in mammalian phased assemblies due to the relatively lower heterozygosity level and the way it is distributed along the genome.

**Figure 2.**
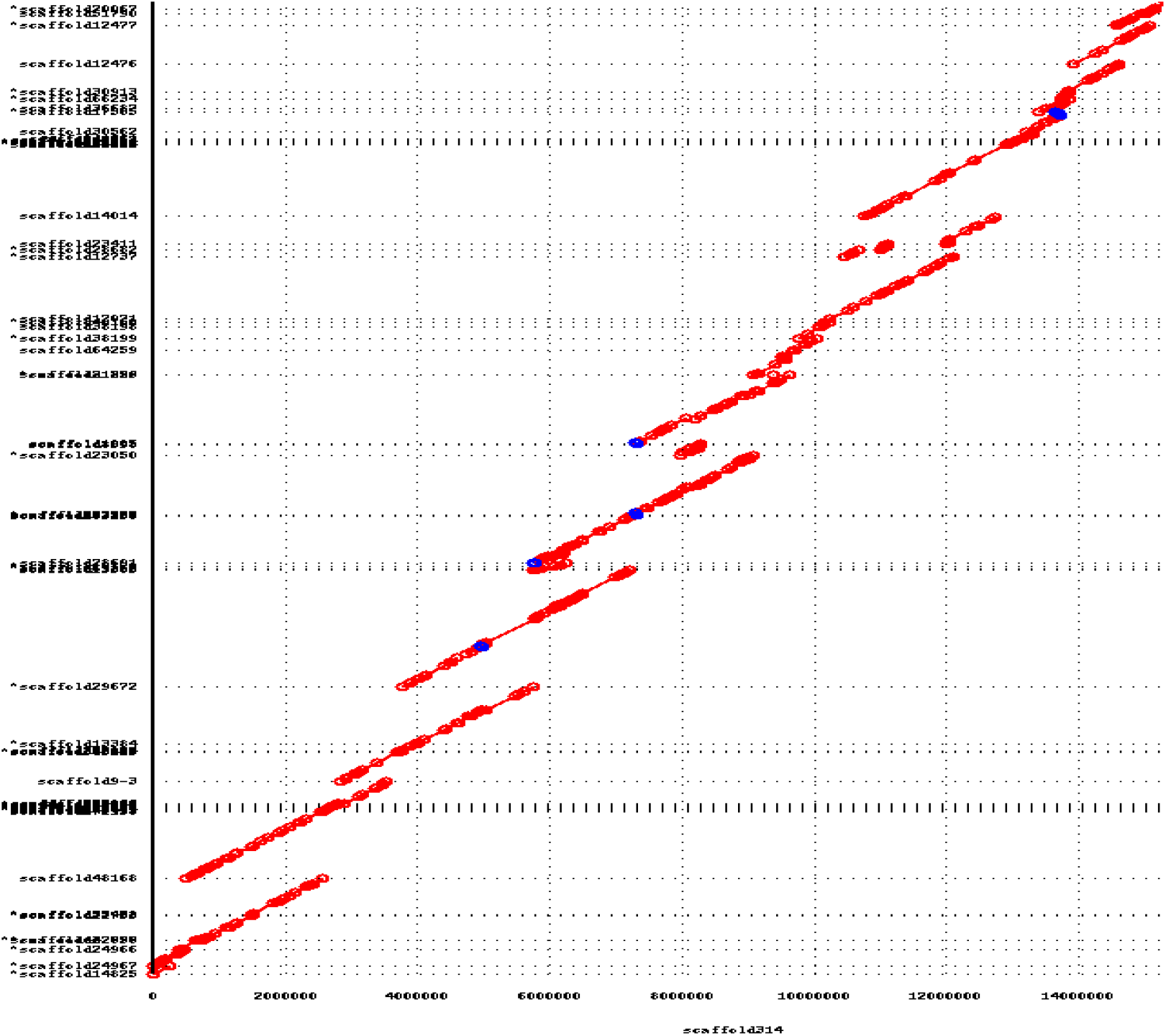
An example mummerplot of the alignments of the phased assembly to the “haploid” one, spanning about 15Mbp of sequence of scaffold314. The circles represent contig ends with lines joining them representing aligned sequence. The color indicates direction of the alignment with red and blue forward and reverse respectively. We show that for most locations on the x-axis (haploid assembly coordinate) there are two alignments on the y-axis corresponding to the two phased haplotypes. The small number of regions with a single contig aligning represent long homozygous regions of the genome that we were unable to phase.

In haplotype phasing it is easy to phase small regions. For example, a single isolated SNP with no haplotype differences within 100 bp in both directions, can be trivially phased into two 201bp (or longer) contigs different by one base in the middle. It gets more difficult for larger contigs/scaffolds, where one must make sure that the contig/scaffold represents single haplotype and not a “mosaic” of haplotypes, that the SNPs and other bigger haplotype differences are correctly “phased”. To do that we mapped the mate pairs from the 5-7Kb mate pair library to all phased scaffolds using Bowtie2 [28], and then examined the “internal” mate pairs where both reads in each pair mapped to the assembly, and one read mapped within 10Kb away from the ends of the scaffold. This would imply that the other mate must map to the same scaffold and not its haplotype, if haplotype phasing is done properly. If it does not, then it indicates an apparent mis-assembly or failure to phase haplotypes. By measuring the number of “properly” aligned internal mates, where both mates aligned to the same scaffold vs. “improper” internal mates where the mates aligned to different scaffolds, one can measure the efficacy of the haplotype phasing. There were 35,697,369 pairs where both mates mapped properly to the same scaffold, while only 169,244 mapped improperly, that is to two different scaffolds. The percentage of improperly mapping mate pairs is only 0.5%, indicating that haplotype resolution was done properly.

### Synteny between human and dolphin

Dolphin is a mammal, and currently the best mammalian reference genome is the human genome. To understand similarities between dolphin and human on the DNA level, we aligned the Tur_tru_Illumina_hap_v1 assembly to the primary chromosomes of the current haploid human reference genome GRCh38 [26]. Since human and dolphin are fairly distant species, we did not expect to find long DNA sequence alignments but instead we were looking for synteny where relatively short DNA fragments of scaffolds align in the same order and orientation between the two assemblies. We used MUMmer4 package for producing the alignments using the default settings. The alignment mummerplot (Figure 3) shows a striking synteny between the dolphin assembled scaffolds and human chromosome 1, visible even on the large-scale chromosome plot (Figure 3a). No large-scale synteny to the other human chromosomes can be readily observed. The synteny observation is possible due to large scaffold sizes in Tur_tru_Illumina_hap_v1. In Figure 3b, we show 22 scaffolds that have 50% or more of their sequence in syntenic alignments. The syntenic alignments of these 22 scaffolds span nearly the entire human chromosome 1 sequence. The synteny is not a new finding, it was first identified by Bielec et al [22] and was later extended to many other placental mammals [23]. The Tur_tru_Illumina_hap_v1 assembly clearly illustrates and confirms the expected synteny.

**Figure 3.**
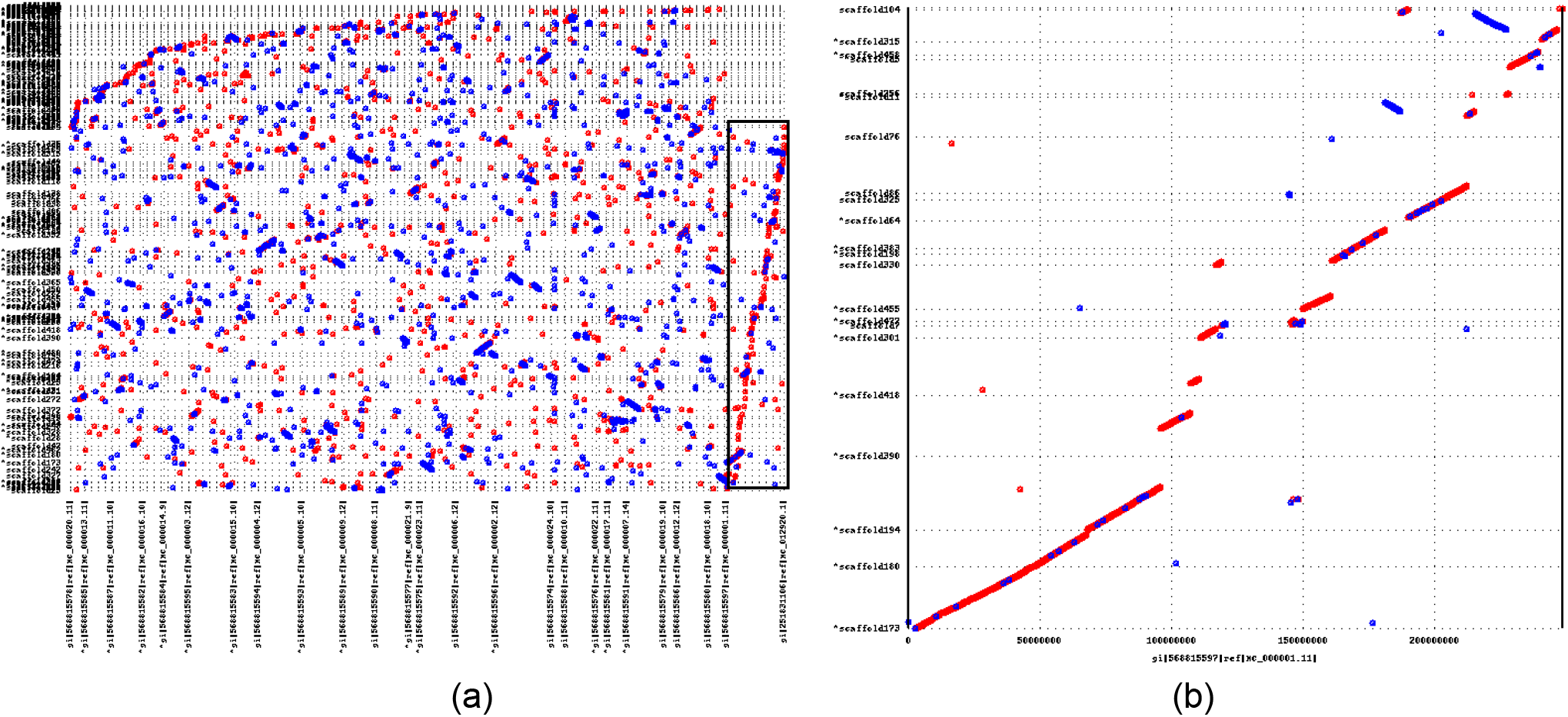
This figure shows the alignment of the Tur_tru_Illumina_hap_v1 assembly to the human GRCH38 reference (primary chromosomes only). Each dot represents an alignment with red indicating forward direction and blue indicating reverse direction. Human reference coordinates are on the X axis and Tur_tru_Illumina_hap_v1 assembly alignment coordinates are on the Y axis. Panel (a) shows alignment of the entire assembly to the human reference with alignments to human chromosome 1 highlighted by the black box. One can clearly see the synteny that is present between the dolphin scaffolds and human chromosome 1. No other human chromosome shows clear synteny. Dolphin scaffolds with syntenic alignments spanning over 50% of the scaffold were extracted. Alignments only to human chromosome 1 are presented in panel (b).

## Methods

### Sample Collection and DNA Extraction

The sample for this study came from a female Atlantic bottlenose dolphin (Sample ID 04329), captive born at SeaWorld of Orlando, Orlando, Florida from wild male and female Atlantic bottlenose dolphins. The animal was 36 years old at blood collection with a healthy medical history. Blood was collected using Qiagen-PAXgene^™^ Blood DNA Tubes (Qiagen). High molecular weight genomic DNA was isolated using the Illumina-MasterPure™ DNA Purification Kit and subsequently quantified and qualified using Quant-iT^™^ dsDNA Kit and E-Gel^™^ EX Agarose Gel (ThermoFisher).

### Collecting Sequence Data

#### Paired End (PE) libraries

We generated the 450bp and 800bp PE libraries using the Illumina TruSeq® PCR-free DNA Sample Prep kit. The protocol was slightly modified at fragmentation and double-size selection steps by adjusting the Covaris DNA shearing protocols and by empirically titrating the ratios of SPRI magnetic beads over DNA to obtain insert sizes around 450bp and 800bp. We then evaluated the libraries for insert size and yield using Agilent Bioanalyzer and real-time qPCR assay, using Illumina DNA Standards and primer master mix qPCR kit (KAPA Biosystems, Roche), then normalized to 2nM prior to clustering and sequencing. Both the 450bp and 800 bp libraries were then denatured and diluted to 8pM and 12pM respectively. The 800bp PE library was clustered and sequenced on the HiSeq 2000, using the Illumina HiSeq Cluster and SBS v4 kits for PE 2 × 160bp reads. The 450bp PE library was clustered and sequenced on the HiSeq 2500 v2 Rapid Run mode using the HiSeq Rapid Cluster and SBS v2 kits for PE 2 × 250bp reads.

#### Mate Pair (MP) libraries

To maximize sequence diversity and genome coverage, three separate MP libraries were constructed corresponding to 2-5Kb, 5-7Kb and 7-10Kb insert sizes using the Nextera® MP Library Preparation Kit according to the manufacturer’s instructions (Illumina). All three libraries were generated from a single input of 4ug of genomic DNA size-selected on a 0.8% E-gel (Invitrogen). Proper sizing of gel-extracted products was confirmed using the Bioanalyzer High Sensitivity chip (Agilent) and 600ng was subsequently used as input for circularization. Following library preparation, the Bioanalyzer was used to confirm library quality. Each of the three libraries were quantified by qPCR (KAPA Biosystems Library Quantification Kit, Roche), denatured and diluted to 200pM after size-adjustment according to Bioanalyzer results, and clustered on the cBot (Illumina) according to the manufacturer’s instructions. 2 × 150bp of Illumina paired-end sequencing was performed on the HiSeq 4000 using the HiSeq 3000/4000 Cluster and SBS kits.

#### 10x Chromium library

Genomic DNA quality was assessed by pulsed-field gel electrophoresis to determine suitability for 10X Chromium library preparation (10X Genomics). 1.125ng of input was used for library preparation according to the manufacturer’s instructions without size-selection. Final library concentration was determined by qPCR (KAPA Library Quantification Kit, Roche) and size-adjusted according to Bioanalyzer DNA 100 chip (Agilent) results. 2 × 150bp of Illumina paired-end sequencing with an 8-base index read was performed on the HiSeq 4000 using the HiSeq 3000/4000 Cluster and SBS kits.

#### Genome Assembly

Genome assembly was conducted using the DeNovoMAGIC^TM^ software platform (NRGene, Nes Ziona, Israel). This is a proprietary DeBruijn-graph-based assembler that was used to produce assemblies of several challenging plant genomes such as corn [1] and ancestral wheat Aegilops tauschii [3]. The following outlines design of the assembler, and steps of the assembly process.

#### Reads pre-processing

In the pre-processing step we first removed PCR duplicate reads, and trimmed Illumina adaptor AGATCGGAAGAGC and Nextera® linker (for MP library) sequences. We then merged the PE 450bp 2×250bp overlapping reads with minimal required overlap of 10bp to create stitched reads (SRs) using the approach similar to the one implemented in the Flash software [27].

#### Error correction

We scanned through all merged reads to detect and filter out reads with apparent sequencing errors by examining k-mers (k=24) in the reads and looking for low abundance k-mers. We have high coverage data (~450x), with each read yielding 127 (150-24+1) to 227 (250-24+1) K-mers. Thus average 24-mer coverage is at least 300x. 24-mers that only appear less than 10 times in the set of reads likely contain errors. We did not use the reads that contain these low abundance k-mers for building initial contigs.

#### Contig assembly

The first step of the assembly consists of building a De Bruijn graph (kmer=127 bp) of contigs from all filtered reads. Next, paired end and MP reads are used to find reliable paths in the graph between contigs for repeat resolving and contigs extension. 10x barcoded reads were mapped to contigs to ensure that adjacent contigs were connected only when there is evidence that those contigs originate from a single stretch of genomic sequence (reads from the same two or more barcodes were mapped to the same contigs).

#### Split phased/un-phased assembly processes

Two parallel assemblies take place to complete the phased and un-phased assembly result. The phased assembly process utilizes the complete set of contigs. In the un-phased assembly process, the homologous contigs are identified and one of the homologs is filtered out, leaving a subset of the homozygous and one of the homologous contigs in heterozygous regions. The linking information of both homologous contigs is kept through the assembly process of the un-phased assembly, usually enabling longer un-phased scaffolds.

#### Scaffolding

All the following steps are done in parallel for both the phased and un-phased assemblies. Contigs were linked into scaffolds with PE and MP information, estimating gaps between the contigs according to the distance of PE and MP links. In addition, for the phased assembly, 10x data were used to validate and support correct phasing during scaffolding.

#### Gap filling

A final gap fill step used PE and MP links and De Bruijn graph information to locally construct a unique path through the graph connecting the gap edges. The path was used to close the gap if it was unique and its length was consistent with the gap size estimate.

#### Scaffold split/merge

We used 10x barcoded reads to refine and merge scaffolds. All barcoded 10x reads were mapped to the assembled scaffolds. Clusters of reads with the same barcode mapped to adjacent contigs in the scaffolds were identified to be part of a single long molecule. Next, each scaffold was scanned with a 20kb length window to ensure that the number of distinct clusters that cover the entire window (indicating a support for this 20kb connection by several long molecules) is statistically significant with respect to the number of clusters that span the left and the right edge of the window. If there was a statistically significant disagreement in the coverage by the clusters over the window, we broke the scaffold at the two edges of the window. Finally, the barcodes that were mapped to the scaffold edges (first and last 20kb sequences) were compared to generate a graph of scaffolds. The scaffolds are nodes and the edges are links connecting nodes with more than two common barcodes on the ends. We broke the links to the nodes that had more than two links and output the resulting linear paths in the scaffold graph as final scaffolds.

## Summary

We show that Tur_tru_Illumina_hap_v1 is more complete and accurate compared to the current best reference Tur_tru v1, based on the amount and composition of sequence, the consistency of the MP alignments to the assembled scaffolds, and on the analysis of conserved single-copy mammalian orthologs. The additional 12.5% of sequence data identified and assembled here was found to contain 165 additional BUSCO alignments as compared to the latest published assembly Tur_tru v1. The large scaffolds represented by Tur_tru_Illumina_hap_v1 enabled and confirmed expected synteny to human chromsome 1. The phased de novo assembly Tur_tru_Illumina_phased_v1 is of the first publicly available and it provides the community with novel ways to explore the heterozygous nature of the dolphin genome. These findings illustrate the impact of improved sample preparation and improved de novo assembly methods on progress toward more complete and accurate reference quality genomes. Better quality assemblies will improve our understanding of gene structure, function and evolution in mammalian species.

## Availability of da

The dolphin assembly Tur_tru_Illumina_hap_v1 has been deposited at NCBI under BioProject PRJNA476133, accession QMGA00000000. The dolphin assembly Tur_tru_Illumina_phased_v1 has been deposited at NCBI under BioProject PRJNA478376, accession QUXD00000000. Both assemblies are also available on the public FTP site ftp://ftp.ccb.jhu.edu/pub/alekseyz/Tur_tru_Illumina_v1/.

## Author contributions

KVM, CTL and MMV designed the project. KVM, CTL and AZ wrote the manuscript. GBZ, TB and KB generated genome assemblies. AZ conducted validation, MP consistency analysis, human chromosome 1 and BUSCO analyses, and submitted the genomes to NCBI; JSL provided the blood sample; JL, AN, MR, MG, EJ, and BS processed samples, generated sequencing and completed quality checks on sequence data; and all authors contributed to editing the manuscript.

## Acknowledgement

This project was sponsored by Illumina, Inc. and NRGene. The authors acknowledge the support of the following people: Tristan Orpin and Matt Posard previously at Illumina, Inc.; Mark Van Oene, Feng Chen, Christine Ching, Shu Boles, Zheng Xu, Joey Flores, Kan Nobuta, Brad Sickler, Courtney McCormick, Christopher Haynes, Jennifer Bernet, and Nathalie Mouttham from Illumina, Inc.; Melissa Katigbak and Shara Fisler from Ocean Discovery Institute; Todd Schmitt from SeaWorld; Edwin Hauw and John Stuelpnagel from 10x Genomics.

## References

1. Hirsch CN, Hirsch CD, Brohammer AB et al. Draft Assembly of Elite Inbred Line PH207 Provides Insights into Genomic and Transcriptome Diversity in Maize. Plant Cell 2016;28:2700–2714

2. Avni R, Nave M, Barad O et al. Wild emmer genome architecture and diversity elucidate wheat evolution and domestication. Science 2017;357:93–97

3. Luo MC, Gu YQ, Puiu D et al. Genome sequence of the progenitor of the wheat D genome Aegilops tauschii. Nature 2017;551:498–502

4. Zimin A, Stevens KA, Crepeau MW et al. An improved assembly of the loblolly pine mega-genome using long-read single-molecule sequencing. GIGAScience 2017;6:1–4

5. Genome 10K Community of Scientists. Genome 10K: A Proposal to Obtain Whole-Genome Sequence for 10000 Vertebrate Species. Journal of Heredity 2009;100(6):659–674

6. Koepfli K, Paten B, Antunes A et al. The Genome 10K Project: A Way Forward Further. Annual Review of Animal Biosciences 2015;3:57–111

7. Lewin HA, Robinson GE, Kress WJ et al. Earth BioGenome Project: Sequencing life for the future of life. Proceedings of the National Academy of Sciences 2018;115(17):4325–4333

8. Mohr DW, Naguib A, Weisenfeld N et al. Improved *de novo* Genome Assembly: Linked-Read Sequencing Combined with Optical Mapping Produce a High Quality Mammalian Genome at Relatively Low Cost. bioRxiv 2017;128348

9. Armstrong EE, Taylor RW, Prost S et al. Entering the era of conservation genomics: Cost-effective assembly of the African wild dog genome using linked long reads. bioRxiv 2017;195180

10. Hammond PS, Bearzi G, Bjørge A et al. *Tursiops truncatus*. The IUCN Red List of Threatened Species 2012:e.T22563A17347397. http://dx.doi.org/10.2305/IUCN.UK.2012.RLTS.T22563A17347397.en.

11. Rosel PE, Hancock-Hanser BL, Archer FI et al. Examining metrics and magnitudes of molecular genetic differentiation used to delimit cetacean subspecies based on mitochondrial DNA control region sequence. Special Issue: Delimiting subspecies using primarily genetic data. Marine Mammal Science 2017;33(S1):76–100

12. McGowen MR, Grossman LI, Wildman DE. Dolphin genome provides evidence for adaptive evolution of nervous system genes and a molecular rate slowdown. Proc. R. Soc. B 2012;279:3643–3651

13. Venn-Watson S, Carlin K, Ridgway S. Dolphins as animal models for type 2 diabetes: sustained, post-prandial hyperglycemia and hyperinsulinemia. Gen Comp Endocrinol 2011; 170(1):193–9

14. Venn-Watson S, Smith CR, Stevenson S et al. 0Blood-Based Indicators of Insulin Resistance and Metabolic Syndrome in Bottlenose Dolphins (*Tursiops truncatus*). Front Endocrinol (Lausanne) 2013;4:136

15. Venn-Watson S. Dolphins as animal models for type 2 diabetes: sustained, post-prandial hyperglycemia and hyperinsulineia. Frontiers in Endocrinology 2014; 227

16. Neely BA, Debra L. Ellisor DL et al. Proteomics as a metrological tool to evaluate genome annotation accuracy following de novo genome assembly: a case study using the Atlantic bottlenose dolphin (*Tursiops truncatus*). bioRxiv 2018;254250

17. Sobolesky P, Parry C, Boxall B et al. Proteomic analysis of non-depleted serum proteins from bottlenose dolphins uncovers a high vanin-1 phenotype. Scientific reports 2016;26(6):33879

18. Venn-Watson S, Smith CR, Gomez F et al. Physiology of aging among healthy, older bottlenose dolphins (*Tursiops truncatus*): comparisons with aging humans. J Comp Physiol B. 2011;181(5):667–80

19. Lindblad-Toh K, Garber M, Zuk O et al. A high-resolution map of human evolutionary constraint using 29 mammals. Nature 2011;478:10530

20. Foote AD, Liu Y, Thomas GWC et al. Convergent evolution of the genomes of marine mammals.2014. Nature Genetics 2014;47,3: 272–275

21. Marks P, Garcia S, Alvaro Martinez A et al. Resolving the Full Spectrum of Human Genome Variation using Linked-Reads. bioRxiv 2018;230946

22. Bielec PE, Gallagher DS, Womack JE et al. Homologies between human and dolphin chromosomes detected by heterologous chromosome painting. Cytogenetic and Genome Research 1998;81(1):18–25

23. Murphy WJ, Frönicke L, O’Brien SJ et al. The origin of human chromosome 1 and its homologs in placental mammals. Genome research. 2003;13(8):1880–8

24. Marçais G, Delcher AL, Phillippy AM et al. MUMmer4: A fast and versatile genome alignment system. PLOS 2018;1005944

25. Putnam NH, O’Connell BL, Stites JC et al. Chromosome-scale shotgun assembly using an in vitro method for long-range linkage. Genome Research 2016;26(3):342–50

26. Schneider VA, Graves-Lindsay T, Howe K et al. Evaluation of GRCh38 and de novo haploid genome assemblies demonstrates the enduring quality of the reference assembly. Genome research 2017;27(5):849–64

27. Magoč T, Salzberg SL. FLASH: fast length adjustment of short reads to improve genome assemblies. Bioinformatics 2011;27(21):2957–63

28. Langmead B, Salzberg SL. Fast gapped-read alignment with Bowtie 2. Nature methods. 2012;9(4):357–359. doi:10.1038/nmeth.1923.

